# An Information Theory Framework for Movement Path Segmentation and Analysis

**DOI:** 10.1101/2024.08.02.606194

**Authors:** Varun Sethi, Orr Spiegel, Richard Salter, Shlomo Cain, Sivan Toledo, Wayne M. Getz

**Affiliations:** Department Environmental Science, Policy and Management, University of California, Berkeley, 94720, CA, USA; School of Zoology, Faculty of Life Sciences, Tel Aviv University, Tel Aviv 69978, Israel; Numerus Inc., 850 Iron Point Road, Folsom, CA 95630, USA; Department of Computer Science, Oberlin College, OH 44074, USA; Blavatnik School of Computer Science, Tel Aviv University, Tel Aviv 69978, Israel; School of Mathematics, Statistics & Computer Science, University of KwaZulu-Natal, South Africa

**Keywords:** path segmentation, movement modes, cluster analysis, machine learning, statistical movement elements (StaMEs), canonical activity modes (CAMs), Jensen-Shannon, barn owl (*Tyto alba*)

## Abstract

Improved animal tracking technologies provide opportunities for novel segmentation of movement tracks/paths into behavioral activity modes (BAMs) critical to understanding the ecology of individuals and the functioning of ecosystems. Current BAM segmentation includes biological change point analyses and hidden Markov models. Here we use an elemental approach to segmenting tracks into *µ*-step-long “base segments” and *m*-base-segment-long “words.” These are respectively clustered into *n* statistical movement elements (StaMEs) and *k* “raw” canonical activity modes (CAMs). Once the words are coded using *m* extracted StaME symbols, those encoded by the same string of symbols, after a rectification processes has been implemented to minimize misassigned words, are identified with particular “rectified” CAM types. The percent of reassignment errors, along with information theory measures, are used to compare the efficiencies of coding both simulated and empirical barn owl data for a selection of parameter values and approaches to clustering.

## INTRODUCTION

Studies of how animals move over landscapes provide us with a lens on the social and behavioral ecology of individuals ^1,2^ and a probe on how climate and landscape changes affect the conservation of endangered species ^3–5^. The movement of animals, in the absence of environmental data, is most often represented by time series of relocation data as sequential points in a 2-d Euclidean space. This time series is typically referred to in the biological literature as a “track” or “path” ^6^ and in the mathematical literature as a “stochastic walk” ^7^. Stochastic walks are often treated as generated by one type of movement process, though animal tracks typically cover a mixture of movement modes that, at a subdiel scale, may include resting, commuting, searching, and foraging ^8,9^), and at a finer scale, bouts of standing, stepping, trotting, jumping etc ^10^. Identifying distinct movement phases is critical for linking observed movement with the underlying ecological process (e.g., predation escape events ^11^). Thus, a key concept in the analysis of animal movement tracks is path segmentation ^6,12^. Its methods focus on detecting changes in movement behavior (e.g.: behavioral change point analyses—BCPA ^13–17^; and hidden Markov models—HMM ^18–22^), also referred to as movement modes ^8,23,24^. These methods work well at the time scale of sub-hours, hours ^25^ and days ^26–28^, but become challenging to apply at finer time scales of minutes and sub-minutes because of, firstly, the need to collect data at frequencies measured in seconds and, secondly, the computational challenges of working with large sets of data.

An alternative to BCPA and HMM approaches to path segmentation is to view relocation time series as a concatenated string of basic movement elements. Unfortunately, as discussed elsewhere ^29^, a set of fundamental movement elements (FuMEs ^30,31^)—such as one stride, one gallop, or one wing flap—is, in general, not possible to identify purely from relocation times series data. One can, however, define the statistical properties of the displacements, average velocities, and turning angles of short base or “symbol” length segments of relocation time series data ^32–34^ and then try to categorize these base segments using clustering methods ^35,36^, including machine learning approaches ^37–39^.

Once we have a set of symbols as our base (or “letter”) level coding segments, they can then be used to construct next level “word” segments of different types ^29,31^. For example, representative symbol segments, computed as the centroids of different clusters, can be used to define a set of statistical movement elements (StaMEs; e.g., large steps with little turning, intermediate steps with moderate turning, and small steps with random changes in direction—see ^29^) that in turn provide a basis for constructing larger fixed-step-number canonical activity mode (CAMs) elements ^30,31^ (e.g., fast persistent movement, meandering movement, resting). Homogeneous strings of CAMs or characteristic heterogeneous mixes of CAMs can then used to define variable length behavioral movement modes (BAMs; e.g., bee lining, searching, foraging) that are currently identified using BCPA or HMM methods, and even larger fixed-time-period diel activity routines (DARs, ^31,40^), such as central placed foraging or nomadism ^41^.

The utility of the methods presented in this paper are manifold and can be used to explore a number of different ecological issues. First, one can use the various measures associated with our method to assess how well organized movement patterns are compared to random patterns constructed out of the same StaME building blocks (i.e., set of symbols). Second, our method provides a way to address hypotheses on how the information content of tracks is affected by an individuals experience in a particular area, an individual’s age, sex, personality type, or state of health, as well as physical and social environmental factors such as season, topography, drought, conditions, density of conspecifics, and presences of competitors, predators, pathogens and parasites. Third, one can asses how movement patterns are shaped as a function of environmental factors and use this information to predict how the movement behavior of an individual may respond to changes in the landscape and, ultimately, climate.

### OUTLINE

Numerical experiments in generating a set of StaMEs from movement track data has been undertaken elsewhere ^29^, though not experiments in generating a set of CAMs and coding these CAMs using the generated StaMEs. This latter dual generating and coding process involves a number of ad-hoc, but considered, decisions (see Tasks 1-4 in Box 1). These include (Fig 1) selecting: i) a value *µ* for the number of consecutive steps used per base (symbol/letter) segment to parse the movement track in a continuous string; ii) the number of categories *n* into which these base segments are sorted and then identified with one of *n* StaME types or symbols (which are centroids of base-length segment clusters); iii) the number *m* of consecutive base segment elements used per word segment (i.e, each word is *mµ* steps long) to parse the movement track in a segmented word string; iv) the number of categories *k* into which these word segments are sorted and then identified with one of *k* raw CAM types (which are centroids of word clusters).

**FIGURE 1.**
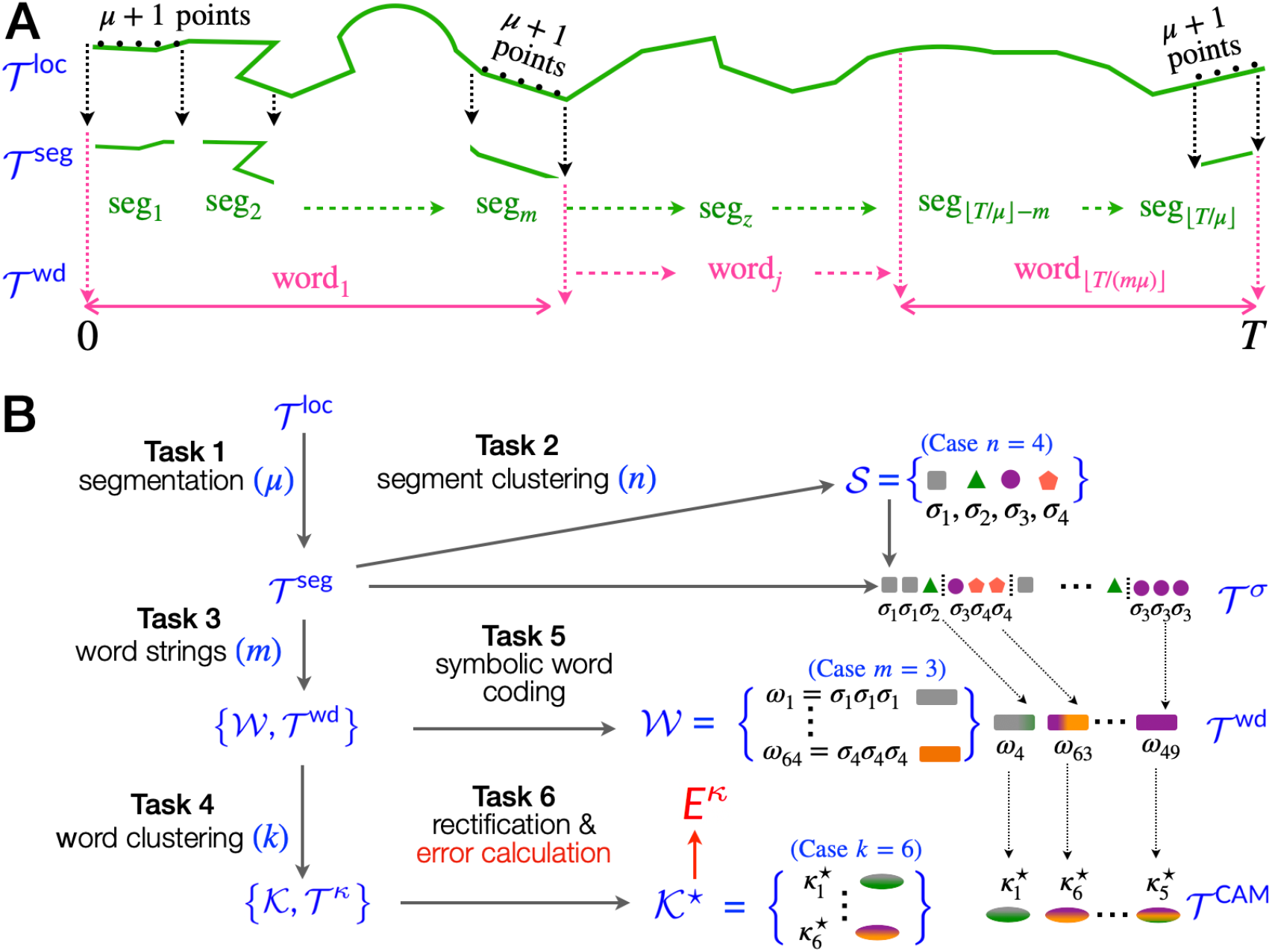
**A**. A graphic illustration of the StaME (clustering of seg_*z*_ elements) and CAM (clustering of word*j* elements) segmentations of the *T* + 1-point relocation data time series *T* ^loc^ (Task 1, Box 1) into the [*T*/*µ*]-segment time series *T* ^seg^ (each segment consists of *µ* steps) and the [*T*/(*µm*)]-word times series *T* ^wd^ (each word consists of *m* segments). **B**. A flow chart of the steps taken, as detailed in BOX 1, to produce segment, StaME (*S*), word, as well as raw (*K*) and rectified (*K*^⋆^) CAM elements. These elements are used to produce segment and word time series *T* ^seg^ and *T* ^wd^, as well as a StaME coded time series *T* ^*σ*^ that is aggregated into a raw CAM time series *T* ^*κ*^ that with reassignments is used to obtain a rectified time series *T* ^CAM^ and an associated assignment error rate E^*κ*^. For the particular case illustrated (*n* = 4, *m* = 3, *k* = 6) the word numbering (*l* = 1, …, 64) scheme is given by Eq 12.

The formulaic underpinnings of the method presented here for segmenting a movement relocation time series *T* ^loc^ into a StaME-coded time series *T*^*σ*^ and a CAM-code time series *T* ^*κ*^ are outlined in ^42^, but numerically evaluated here using both simulated and empirical owl (*Tyto alba* ^29,40,43^) data. We begin our presentation with a quick review of hierarchical information coding concepts ^44^, including the measures we use to compare the efficiency and information content of coding relocation data track into StaME (*T* ^*σ*^) and CAM (*T* ^*κ*^) time series as a function of parameters *µ, n, m*, and *k* and methods of clustering. In Box 1 (also see Fig 1), we summarize the 6 tasks needed complete the coding process. We also briefly discuss salient aspects of the clustering methods used in our numerical experiments with most of our discussing, including references to the broader clustering methodology literature ^9,35,39,45,46^ appearing in Appendix B (SOF).

### HIERARCHICAL CODING

The material presented in this section and Box 1 was conceptualized in ^42^, though without any computational development of the methodology presented below. Here we provide the reader with sufficient details to appreciate the source and derivation of the measures used to compare the different methods employed for parsing the tracks of both simulation and ban owl movement data—i.e., 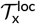 for x=sim and owl— into StaME 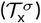, raw CAM 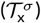, and rectified CAM 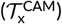 tracks, following the approach described in Box 1, Tasks 1-6.

The amount of information that is coded into the track *T* ^*σ*^ by the symbols *σ*_*i*_ ∈ *S*, when each occurs with uncorrelated probabilities **p**^*σ*^ (Task 2, Box 1), according to Shannon Information Theory ^47,48^, is given by the information entropy measure

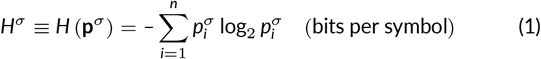

Similarly, the amount of information in the raw and rectified CAM strings *T* ^*κ*^ and *T* ^CAM^ is

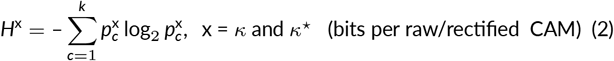

The maximum number of bits of information carried by each symbol and each CAM in describing the trajectory at their respective levels of analysis (*µ* steps per StaME, *mµ* steps per CAM) occurs when all elements are equally likely. This implies

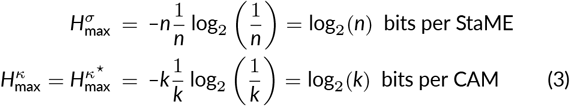

which leads us to define the efficiency of a particular coding scheme

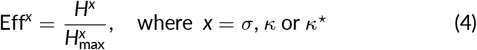

If the proportion of *ω*_*l*_ word types in *W*_*c*_, derived when implementing Task 3 (Box 1), is given by the vector 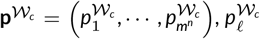 for *l* = 1, …, *n*^*m*^ then we can use the following normalized (to a percentage) Jensen-Shannon divergence measure ^44^ across the ensemble of discrete distributions **p**_*c*_, each of sample size 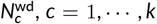 with respect to its mixture distribution, which is just the distribution of word types **p**^**W**^ in the original word set *W* ^42^:

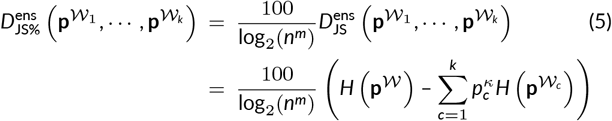

Thus the Jensen-Shannon ensemble divergence measure is the amount of information gained (i.e., reduction in log base 2 entropy) by clustering the word set *W* into the *k* groups *W*^*c*^. The centroids (within group averages of the statistics of all elements in each cluster) provide the raw CAM set *K*.

As indicated in Box 1, the method relies on the selection of the 4 parameters, *µ* (base segment size), *n* (number of segment clusters used to derive the StaME set *S*), *m* (number of base segments in a word), *k* (number of word clusters used to derive the CAM set *K*) and on both base segment and word clustering algorithms denoted by CM^seg^ and CM^wd^. Thus our general method ℳ, in terms of free parameter and clustering method arguments, can be expressed as the functor

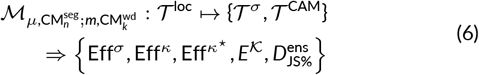

## DATA TREATMENT AND ANALYSES

In this section, we formalize the method used to construct sets of symbols and words from relocation data and apply it to a set of simulated movement data, 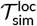, derived from the simulation models Numerus AN-IMOVER_1^29^, and a barn owl (*Tyto alba*) movement track, 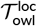, obtained using an ATLAS reverse GPS installation ^49,50^ in the Harod Valley in Israel ^29,43^.

### BOX 1

**Segments, Clusters & Codes** (Fig 1)

**Task 1**. Parse the relocation data series

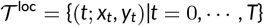

into *µ*-step “base” segments to obtain the data series

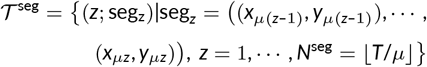

**Task 2**. Cluster base segments in *T* ^seg^ in the *n* StaME set of symbols

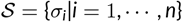

with frequency distribution vector 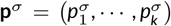 of symbols in

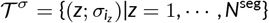

**Task 3**. Parse *T* ^loc^ = {(*t*; *x*_*t*_, *y*_*t*_)|*t* = 0, …, *T*} into (*mµ*)-step “word” segments to obtain the data series

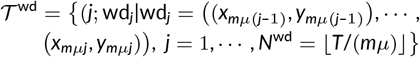

**Task 4**. Cluster word segments in *T* ^wd^ into a set of *k* word clusters *W*_*c*_ each containing 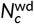 words, *c* = 1, …, *k*. Use cluster centroids *κ*_*c*_ (obtained by averaging across the vectors Eq 9 of all segments in the same cluster) to code *T* ^wd^ into the “raw” CAM sequence and CAM cluster

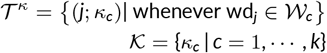

with frequency of occurrence 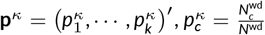

**Task 5**. Code words using StaMEs by comparing the symbolic coding of *T* ^*σ*^ with the word coding of *T* ^wd^ to obtain

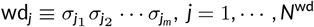

and then identify the 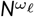 identically coded words with one of the *n*^*m*^ possible types of words *ω*_*l*_ in the set

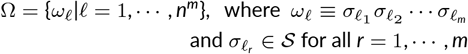

**Task 6**. Rectify the CAM set *K* by identifying the word type *ω*_*l*_ with the CAM *κ*_*c*_ where this word occurs more frequently in **W**_*c*_ than any other **W**_*c*′_ for *c*^*′*^ ≠ *c, c* = 1, …, *k* and transfer the misassigned words to the cluster of its defining type. Denote the set of such rectified CAMs by *K*^⋆^, where the centroid of each of the reorganized clusters is denoted by 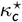: i.e.,

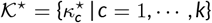

with frequency of occurrence in 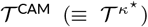 given by 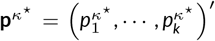. Compute the assignment percentage error

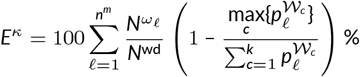

### Relevant variables

Our simulated movement data, denoted 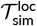, was synthesized using ANIMOVER_1, a two-mode step-selection kernel simulator built using Numerus RAMP technology ^29^. Our barn owl data ^43^, denoted 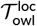, was collected at a relocation frequency of 4 sec and comes from an adult female as reported in ^29^. To begin with, the data was cleaned and re-organized into a shape suitable for our workflow at [VS’s GitHub repository](https://github.com/Observarun/statistical-movement-elements_canonical-activity-modes_CAM-coding-with-StaMEs)

Two primary variables, derived directly from the *T* ^loc^ time series, are the speed (step-size/time-interval) time series and angle of heading time series ^31^

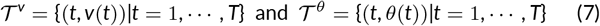

with the latter used to respectively derive the turning angle (change in angle of heading) and absolute turning angle time series (also see Appendix A.2 in ^29^)

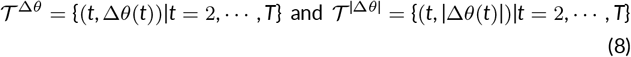

### Clustering methods

The methods we used to cluster the data can be divided into two classes: vector methods that use euclidean and machine learning approaches ^9,35–38^, and shape methods that use deep learning approaches ^39,45,46^. For our vector clustering methods, each element is a base segment, seg_z_, represented by the five variable vector ^29^

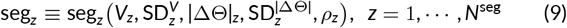

In these vector cases, after categorizing all seg_*z*_ into *n* clusters, the StaMEs *σ*_*i*_ for cluster *i* = 1, …, *n* are taken as the centroid value statistics (i.e., taking averages, denoted by “bar”, across all the same arguments of each seg_*z*_ in cluster *i*) to obtain the vector

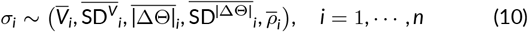

Similarly, the *k* CAMs are taken as the centroids of the *k* word clusters (i.e., taking averages, denoted by “hat”, across all words in cluster *k*) and denoted by

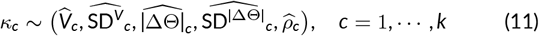

Shape-based clustering methods are applied directly to the time-series data Eq 7 and 8. We used two closely related unsupervised learning methods methods referred to as dynamic time warping, the first using a hard-min and the second a soft-min computational procedure ^51^. Once the clusters were produced by these shape methods, the centroids of these clusters were computed in the same way as for the vector clustering methods: that is, the vector statistics for every segment in the cluster (even thought these statistics were not used in the shape clustering procedure) were averaged across all segments in the cluster to obtain the vector representation of the cluster centroids.

A brief summary of the clustering methods we used—which we refer to as Ward’s Euclidean (E) ^36^, graph spectral (S) ^52^, random forest (F) ^53^, dynamic time warping ^5 4^ (denoted by D and when applied to the absolute values of TA in Eq 8, and by D*′* when applied to the actual values of TA in Eq 8)) and soft-min d ynamic warping ^5 1^ (M)—are provided in Appendix B (SOF).

### StaMEs and Word distributions

After i mplementing th e in itial par t of 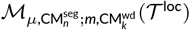 to obtain *T* ^*σ*^ (Task 2, Box 1) and the next part to obtain *T* ^wd^ (Task 2, Box 1), each of the *N*^wd^ words wd_*j*_ ∈ *W* needs to be assigned a word type *ω*_*l*_, *l* = 1, …, *n*^*m*^ (Task 5, Box 1). Various possibilities exist for numbering these types in terms of the sequence of underlying symbols *σ*_*i*_ ∈ *S* defining each type. Here we propose using a sorting algorithm based on ordering the symbols *σ*_*i*_ according to their average velocity 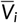 (Eq 10) such that *V*_*i*_ ≥ *V*_*i*+1_ is ensured for all *i* = 1, …, *n* – 1. The word number scheme that we used for the case *n* = 4, *m* = 3, which is relevant to the plots in Fig 2, is:

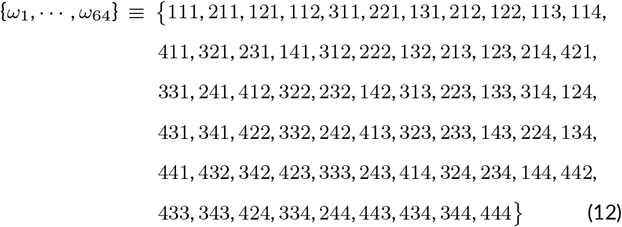

**FIGURE 2.**
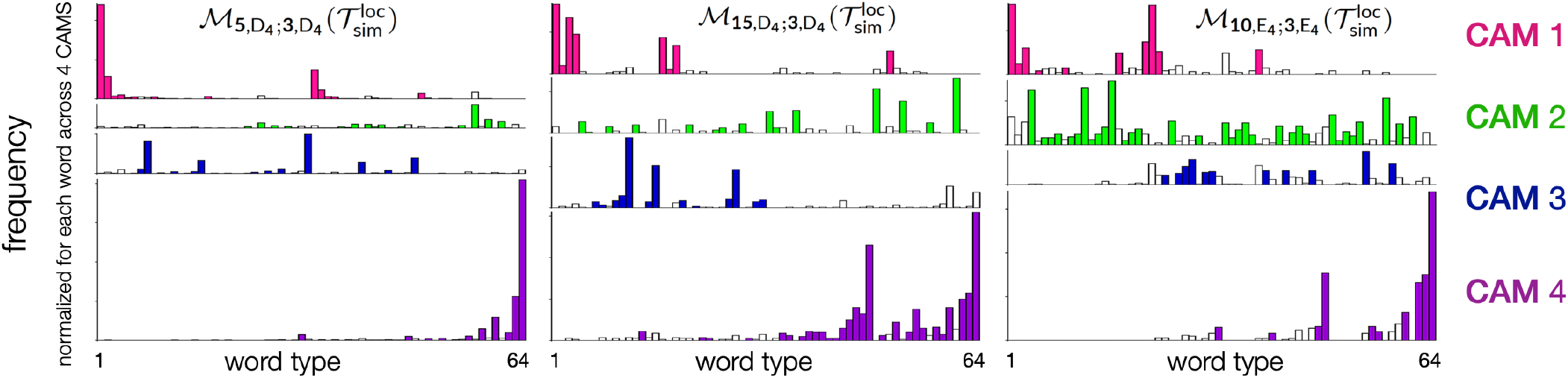
Distribution of word types for 3 of the simulated data cases (Table 3). The bars plotted in each of the 4 ribbons depict the word types (*w*_*l*_, *l* = 1, …, 64 (as labeled according to Eq 12, ordered from fastest to slowest words) for the 4 clusters identified with *κ*_*c*_, *c* = 1, …, 4. Filled columns indicate the assignments of words *ω*_*l*_ to rectified cluster 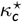, so that white columns are the erroneous assignments used to generate the measure E^*κ*^ percent (Task 6, Box 1).

One of the purposes of the paper is to evaluate the integrity of our StaMEs as the bases for coding rectified CAMs by computing the error E^*κ*^ presented at the bottom of Box 1. In short, after clustering both the base (Task 2, Box 1) and word segments (Task 4, Box 1) of size *µ* and *mµ* steps respectively, the *m* base segments of length *µ* in each raw CAM is replaced with the cluster symbol *σ* (i.e., StaME) to which the particular base segment belongs. Each word is then identified with its StaME type—of which there are *N*^wd^ = 64 that are numbered as depicted in Eq 12. Once this is done, then the distribution of the number of words of each coded type can be plotted for each of the *k* groups into which the words have been clustered (as illustrated in Fig 2 for the case *k* = 4).

Throughout this presentation, we have confined ourselves to the case *k* = 4, which provides sufficient detail for us to demonstrate the utility of approach without an exponential rise in the number of cases that an be presented when results for different values of *k* are compared. We expect, though, that an in-depth application of our methods to a specific group of organisms would need to vary *k* to some extent to obtain the best value for this parameter, which is expected to be somewhat system specific.

Once the set of *k* distributions for the *N*^wd^ words have been obtained, then Task 6 (Box 1) can be implemented to compute the percentage E^*κ*^ of erroneous assignments that occur when rectifying the set of raw CAMs *K* to obtain the “rectified” CAM set *K*^⋆^. This process can be illustrated when plotting the *k* raw CAM distributions of the *N*^wd^ and using closed bars to indicate which of the *k* distributions each of the rectified CAMs *κ*^⋆^ has been assigned to, with open bars representing words that need to be reassigned using the max rule (i.e., assign all words of a particular type to the cluster where the greatest number of them appear—see formula for E^*κ*^ under Task 6 in Box 1), as illustrated in Fig 2 in the next section.

## RESULTS

The results presented here pertain to a simulation data set 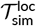 and a barn owl data set 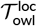 presented elsewhere ^29^. The reader is referred to this study for details on the algorithms used to generate 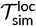. These data consisted of *T* = 69, 527 relocation steps. In this previous study, relocation data sets were clustered into *n* = 8 different types of 10, 15, and 30 point (*µ* = 10, 15 and 30) StaMEs that, if used here to generate CAMs consisting of 3 symbols, would generate *l* = 8^3^ = 512 word types. The word distributions for this case are difficult to visualize graphically, thus we started with a more manageable baseline case of *n* = 4 and *m* = 3, which results in *n*^*m*^ = 64 word types.

### Baseline StaME and CAM Set Results

For illustrative and comparative purposes, we computed a “baseline” set of StaMEs (*S*, Task 2, Box 1) and a raw CAM set (*K*, Task 4, Box 1) for the simulation and owl relocation tracks 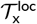, x = sim and owl. For our baseline parameter values we selected *µ* = 10 for the simulation data and *µ* = 15 (i.e., 1 minute segments since consecutive points are 4 min apart) for the owl data. In addition, we set *n* = 4 for the number of distinct types of StaMEs and *m* = 3 for the number of distinct CAMs in both cases. Also, for both the simulation and owl data, our baseline method for clustering base segments into StaMEs and word segments into CAMs was the hierarchical Ward’s approach (E) described in Appendix B (SOF). Thus our baseline methods is

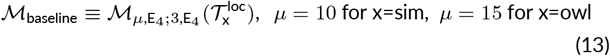

The set of StaME and CAM (Box 1) statistics obtained from the baseline and other methods are reported in Table 1A for the simulation data and 11B for the owl data.

**TABLE 1.** The entries in parts **A** and **B** of this table were obtained using the baseline method presented in Eq 13: they record the centroid statistics and their proportions *p*_*i*_ of occurrence for the StaMEs *σ*_*i*_ (*i* = 1, …, *n* = 4, Eq 10) and raw CAMs *κ*_*c*_ (*c* = 1, …, *k* = 4, Eq 11) of the simulation (**A**, *N*^seg^ = 6, 952) and owl data (**B**, *N*^seg^ = 71, 706) (see Box 1, Tasks 1 and 2). The remaining entries in parts **C**-**E** use the methods to obtain the StaME and now rectified CAM statistics (Box 1, Tasks 5 and 6), as labeled, with the primary difference being the number of StaMEs in the cases considered is *n* = 8, the parameter *m* = 4 rather than 3.

| **A. Baseline simulation: $\mathcal{M}_{10,E_4;3,E_4}(\mathcal{T}_{\text{sim}}^{\text{loc}})$** |  |  |  |  |  |  |
| --- | --- | --- | --- | --- | --- | --- |
|  |  | StaMEs |  |  |  |  |
| $\sigma_i$ | $p_i^\sigma$ | $\bar{V}_i$ | $\overline{SD}_i^V$ | $\vert \overline{\Delta\Theta}\vert _i$ | $\overline{SD}\vert \overline{\Delta\Theta}\vert _i$ | $\bar{p}_i$ |
| $\sigma_1$ | 0.23 | 0.85 | 0.13 | 0.18 | 0.12 | 0.83 |
| $\sigma_2$ | 0.22 | 0.59 | 0.31 | 0.30 | 0.25 | 0.12 |
| $\sigma_3$ | 0.23 | 0.38 | 0.29 | 0.37 | 0.23 | 0.58 |
| $\sigma_4$ | 0.32 | 0.11 | 0.35 | 0.08 | 0.23 | 0.32 |
|  |  | Raw CAMs |  |  |  |  |
| $\kappa_c$ | $p_c^\kappa$ | $\hat{V}_c$ | $\widehat{SD}_c^V$ | $\vert \widehat{\Delta\Theta}\vert _c$ | $\widehat{SD}\vert \widehat{\Delta\Theta}\vert _c$ | $\hat{p}_c$ |
| $\kappa_1$ | 0.19 | 0.77 | 0.22 | 0.29 | 0.21 | 0.20 |
| $\kappa_2$ | 0.40 | 0.48 | 0.25 | 0.38 | 0.22 | 0.62 |
| $\kappa_3$ | 0.16 | 0.39 | 0.31 | 0.39 | 0.25 | 0.14 |
| $\kappa_4$ | 0.25 | 0.18 | 0.34 | 0.21 | 0.23 | 0.17 |

| **B. Baseline owl: $\mathcal{M}_{15,E_4;3,E_4}(\mathcal{T}_{\text{owl}}^{\text{loc}})$** |  |  |  |  |  |  |
| --- | --- | --- | --- | --- | --- | --- |
|  |  | StaMEs |  |  |  |  |
| $\sigma_i$ | $p_i^\sigma$ | $\bar{V}_i$ | $\overline{SD}_i^V$ | $\vert \overline{\Delta\Theta}\vert _i$ | $\overline{SD}\vert \overline{\Delta\Theta}\vert _i$ | $\bar{p}_i$ |
| $\sigma_1$ | 0.02 | 0.066 | 0.039 | 0.21 | 0.22 | 0.89 |
| $\sigma_2$ | 0.08 | 0.015 | 0.013 | 0.36 | 0.30 | 0.52 |
| $\sigma_3$ | 0.46 | 0.012 | 0.008 | 0.39 | 0.32 | 0.09 |
| $\sigma_4$ | 0.43 | 0.011 | 0.007 | 0.39 | 0.31 | 0.23 |
|  |  | Raw CAMs |  |  |  |  |
| $\kappa_c$ | $p_c^\kappa$ | $\hat{V}_c$ | $\widehat{SD}_c^V$ | $\vert \widehat{\Delta\Theta}\vert _c$ | $\widehat{SD}\vert \widehat{\Delta\Theta}\vert _c$ | $\hat{p}_c$ |
| $\kappa_1$ | 0.03 | 0.047 | 0.045 | 0.28 | 0.27 | 0.80 |
| $\kappa_2$ | 0.04 | 0.018 | 0.025 | 0.35 | 0.30 | 0.50 |
| $\kappa_3$ | 0.66 | 0.012 | 0.008 | 0.39 | 0.32 | 0.04 |
| $\kappa_4$ | 0.27 | 0.011 | 0.008 | 0.38 | 0.31 | 0.13 |

| **C. $\mathcal{M}_{10,S_8;5,S_4}(\mathcal{T}_{\text{owl}}^{\text{loc}})$** |  |  |  |  |  |  |
| --- | --- | --- | --- | --- | --- | --- |
|  |  | StaMEs |  |  |  |  |
| $\sigma_i$ | $p_i^\sigma$ | $\bar{V}_i$ | $\overline{SD}_i^V$ | $\vert \overline{\Delta\Theta}\vert _i$ | $\overline{SD}\vert \overline{\Delta\Theta}\vert _i$ | $\bar{p}_i$ |
| $\sigma_1$ | 0.01 | 0.232 | 0.048 | 0.07 | 0.06 | 0.95 |
| $\sigma_2$ | 0.06 | 0.067 | 0.044 | 0.37 | 0.33 | 0.28 |
| $\sigma_3$ | 0.03 | 0.045 | 0.042 | 0.29 | 0.29 | 0.81 |
| $\sigma_4$ | 0.15 | 0.021 | 0.012 | 0.32 | 0.30 | 0.11 |
| $\sigma_5$ | 0.20 | 0.019 | 0.011 | 0.45 | 0.37 | 0.15 |
| $\sigma_6$ | 0.20 | 0.018 | 0.010 | 0.42 | 0.25 | 0.22 |
| $\sigma_7$ | 0.08 | 0.018 | 0.013 | 0.46 | 0.33 | 0.52 |
| $\sigma_8$ | 0.27 | 0.018 | 0.010 | 0.34 | 0.31 | 0.37 |
| | | $S_4$ Rectified CAM cluster | | | | |
| $\kappa_c^*$ | $p_c^{\kappa^*}$ | $\hat{V}_c$ | $\widehat{SD}_c^V$ | $\vert \widehat{\Delta\Theta}\vert _c$ | $\widehat{SD}\vert \widehat{\Delta\Theta}\vert _c$ | $\hat{p}_c$ |
| $\kappa_1^*$ | 0.02 | 0.131 | 0.101 | 0.22 | 0.25 | 0.84 |
| $\kappa_2^*$ | 0.07 | 0.028 | 0.041 | 0.37 | 0.30 | 0.44 |
| $\kappa_3^*$ | 0.47 | 0.024 | 0.017 | 0.40 | 0.33 | 0.05 |
| $\kappa_4^*$ | 0.45 | 0.020 | 0.013 | 0.37 | 0.29 | 0.07 |

| **D. $\mathcal{M}_{10,D_8;5,D_4}(\mathcal{T}_{\text{owl}}^{\text{loc}})$** |  |  |  |  |  |  |
| --- | --- | --- | --- | --- | --- | --- |
|  |  | StaMEs |  |  |  |  |
| $\sigma_i$ | $p_i^\sigma$ | $\bar{V}_i$ | $\overline{SD}_i^V$ | $\vert \overline{\Delta\Theta}\vert _i$ | $\overline{SD}\vert \overline{\Delta\Theta}\vert _i$ | $\bar{p}_i$ |
| $\sigma_1$ | 0.01 | 0.197 | 0.62 | 0.12 | 0.11 | 0.91 |
| $\sigma_2$ | 0.10 | 0.025 | 0.015 | 0.39 | 0.37 | 0.21 |
| $\sigma_3$ | 0.19 | 0.023 | 0.014 | 0.35 | 0.33 | 0.29 |
| $\sigma_4$ | 0.13 | 0.023 | 0.015 | 0.41 | 0.33 | 0.28 |
| $\sigma_5$ | 0.20 | 0.023 | 0.014 | 0.40 | 0.34 | 0.27 |
| $\sigma_6$ | 0.12 | 0.022 | 0.013 | 0.34 | 0.22 | 0.32 |
| $\sigma_7$ | 0.14 | 0.021 | 0.013 | 0.44 | 0.33 | 0.26 |
| $\sigma_8$ | 0.10 | 0.019 | 0.011 | 0.38 | 0.29 | 0.27 |
|  |  | Rectified CAMs |  |  |  |  |
| $\kappa_c^*$ | $p_c^{\kappa^*}$ | $\hat{V}_c$ | $\widehat{SD}_c^V$ | $\vert \widehat{\Delta\Theta}\vert _c$ | $\widehat{SD}\vert \widehat{\Delta\Theta}\vert _c$ | $\hat{p}_c$ |
| $\kappa_1^*$ | 0.005 | 0.198 | 0.091 | 0.13 | 0.18 | 0.81 |
| $\kappa_2^*$ | 0.30 | 0.034 | 0.027 | 0.38 | 0.30 | 0.13 |
| $\kappa_3^*$ | 0.20 | 0.027 | 0.019 | 0.39 | 0.35 | 0.06 |
| $\kappa_4^*$ | 0.495 | 0.015 | 0.011 | 0.38 | 0.30 | 0.09 |

| **E. $\mathcal{M}_{10,D_8';5,D_4'}(\mathcal{T}_{\text{owl}}^{\text{loc}})$** |  |  |  |  |  |  |
| --- | --- | --- | --- | --- | --- | --- |
|  |  | StaMEs |  |  |  |  |
| $\sigma_i$ | $p_i^\sigma$ | $\bar{V}_i$ | $\overline{SD}_i^V$ | $\vert \overline{\Delta\Theta}\vert _i$ | $\overline{SD}\vert \overline{\Delta\Theta}\vert _i$ | $\bar{p}_i$ |
| $\sigma_1$ | 0.01 | 0.188 | 0.066 | -0.01 | 0.20 | 0.87 |
| $\sigma_2$ | 0.23 | 0.023 | 0.015 | -0.19 | 0.44 | 0.27 |
| $\sigma_3$ | 0.14 | 0.022 | 0.014 | -0.08 | 0.48 | 0.27 |
| $\sigma_4$ | 0.15 | 0.022 | 0.014 | -0.04 | 0.48 | 0.26 |
| $\sigma_5$ | 0.09 | 0.022 | 0.013 | 0.29 | 0.38 | 0.26 |
| $\sigma_6$ | 0.14 | 0.021 | 0.013 | 0.08 | 0.51 | 0.27 |
| $\sigma_7$ | 0.13 | 0.021 | 0.013 | 0.11 | 0.51 | 0.27 |
| $\sigma_8$ | 0.10 | 0.020 | 0.013 | 0.05 | 0.51 | 0.29 |
| | | $S_4$ Rectified CAM cluster | | | | |
| $\kappa_c^*$ | $p_c^{\kappa^*}$ | $\hat{V}_c$ | $\widehat{SD}_c^V$ | $\vert \widehat{\Delta\Theta}\vert _c$ | $\widehat{SD}\vert \widehat{\Delta\Theta}\vert _c$ | $\hat{p}_c$ |
| $\kappa_1^*$ | 0.02 | 0.107 | 0.094 | -0.01 | 0.36 | 0.72 |
| $\kappa_2^*$ | 0.18 | 0.029 | 0.021 | -0.04 | 0.49 | 0.08 |
| $\kappa_3^*$ | 0.32 | 0.025 | 0.018 | 0.01 | 0.50 | 0.08 |
| $\kappa_4^*$ | 0.47 | 0.018 | 0.013 | 0.01 | 0.49 | 0.09 |

### Comparative StaME Analysis

We begin our evaluation, by first comparing the characteristics of our five methods, E, S, R, D and M in the context of the measure Eff^*σ*^ (Eq 4, with x=*σ*) when completing the first two tasks in Box 1: i.e., clustering base segments of the simulation data 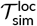 (Eq 10) to generate the StaME set *S*. In particular, we compare these methods using four different base segment step-lengths *µ* = 5, 10, 15 and 30 and two different StaME set sizes *n* = 2 and 4. The results of these 5 × 4 × 2 = 40 different approaches are reported in Table D1 (SOF). All things equal, a higher Eff^*σ*^ score implies a more effective (i.e., informative) coding scheme. However, the distribution of different movement modes might well be skewed in practice (implying a lower Eff^*σ*^ score) because some movements are naturally rarer than others (e.g., occasional fast dashes versus a prevailing normal gait in both predators and prey). Consequently, the measure Eff^*σ*^ on its own does not provide definitive insights into the utility of a particular set of parameters and clustering approaches. Arguably more salient than the measure Eff^*σ*^ may be the span of the velocity values associated with the elements *σ*_*i*_ ∈ *S* encountered under the 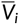 column in the main text Table 1.

The only consistent message that we can take away from the results reported in Table D1 is that the random forest clustering method *F* consistently yield much lower values for Eff^*σ*^ than the other four methods. For example, for the case *µ* = 15 and *n* = 2, the values obtained for E, S, F, D, and M respectively are Eff^*σ*^ = 1.0, 0.97, 0.38, 0.85 and 0.97 (Table D1A). Similarly, for the case *µ* = 10 and *n* = 4, the values obtained for E, S, F, D, and M respectively are Eff^*σ*^ = 0.99, 0.97, 0.55, 0.91 and 0.93 (Table D1B). Given this obvious odd-man-out performance of F, and our need to pare down the number of comparisons we make to a manageable size, we decided to focus on comparison’s primarily between the spectral vector method S and the shape method D (which appears to give similar results to the shape method M). This allows us to focus on a vector versus a shape method comparison. This is not to say that the random forest method F or soft-min method (M) may not prove useful in future studies, the latter for being less computationally demanding than than its twin D method.

### Comparative CAM Analysis

Continuing beyond tasks 1 and 2 (Box 1), once we have completed the remaining tasks 3-6, we can then compare the information coding rates, coding efficiencies, and raw-to-rectified CAM assignment errors for different sets of parameters and clustering methods. In particular, we can now compare the centroid statistics for both the StaMEs and CAMs produced by our method for selected combinations of the following parameter values: *µ* = 5, 10, and 15, *n* = 4 and 8, *m* = 3 and 5, *k* = 4 only, and clustering methods E (for baseline cases only), S and D (including D^*′*^ which uses the actual rather than absolute valued turning angle time series).

Details of the centroid statistics obtained for both the StaME and CAM sets—raw CAMs in Table 1A&B, rectified CAMs in Table 1C-F— and of the distributions of StaMEs in these latter 3 rectified CAM sets (Table 2) are only presented for our most germane cases. The selection of these cases was based on a comparison of the performance measures Eff^*σ*^, Eff^*κ*^, 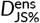 and E^*κ*^ reported in Table 3.

**TABLE 2.** Percent of the 8 StaME types found in each of the four rectified CAMs extracting using indicated methods (Table 2)

| CAM | $\kappa_1^*$ | $\kappa_2^*$ | $\kappa_3^*$ | $\kappa_4^*$ |
| --- | --- | --- | --- | --- |
| | $\mathcal{M}_{10,S_8;5,S_4}$ | | | |
| $\sigma_1$ | 42.8 | 0.10 | 0.0 | 0.1 |
| $\sigma_2$ | 7.0 | 6.7 | 8.2 | 4.1 |
| $\sigma_3$ | 21.4 | 18.1 | 1.2 | 2.0 |
| $\sigma_4$ | 3.5 | 9.8 | 14.7 | 16.7 |
| $\sigma_5$ | 5.1 | 15.3 | 31.8 | 10.1 |
| $\sigma_6$ | 5.8 | 15.9 | 12.1 | 29.0 |
| $\sigma_7$ | 4.0 | 10.0 | 9.0 | 6.3 |
| $\sigma_8$ | 10.5 | 23.3 | 23.0 | 31.8 |
| | $\mathcal{M}_{10,D_8;5,D_4}$ | | | |
| **StaME** |  |  |  |  |
| $\sigma_1$ | 73.6 | 2.3 | 0.1 | 0.1 |
| $\sigma_2$ | 2.1 | 7.0 | 18.2 | 8.4 |
| $\sigma_3$ | 7.1 | 19.4 | 16.5 | 20.1 |
| $\sigma_4$ | 5.7 | 13.8 | 16.3 | 12.3 |
| $\sigma_5$ | 2.1 | 18.1 | 24.2 | 19.2 |
| $\sigma_6$ | 6.4 | 15.0 | 3.4 | 14.2 |
| $\sigma_7$ | 1.4 | 14.1 | 15.2 | 14.2 |
| $\sigma_8$ | 1.4 | 10 | 6.1 | 11.4 |
| | $\mathcal{M}_{10,D'_8;5,D'_4}$ | | | |
| $\sigma_1$ | 41.9 | 0.3 | 0.3 | 0.3 |
| $\sigma_2$ | 18.4 | 31.3 | 21.9 | 21.5 |
| $\sigma_3$ | 9.2 | 14.9 | 12.2 | 14.6 |
| $\sigma_4$ | 8.3 | 15.5 | 16.2 | 14.4 |
| $\sigma_5$ | 4.4 | 6.4 | 10.4 | 9.8 |
| $\sigma_6$ | 6.1 | 13.2 | 13.3 | 15.2 |
| $\sigma_7$ | 5.5 | 10.1 | 14.6 | 13.5 |
| $\sigma_8$ | 6.1 | 8.3 | 11.1 | 10.5 |

**TABLE 3.** For *x* = *σ, κ* and *κ*^⋆^ the percent efficiencies Eff^*x*^ (Eq 4) are listed for the different approaches (index *a* in Eq 6) along with the percent normalized Jensen-Shannon ensemble divergences 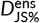 (Eq 5) and the percent assignment errors E^*κ*^ (Task 6, Box 1) with regard the nine approaches used to code the movement track 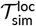 and the 17 approaches used to code the movement track 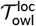. The notation used is that of Eq 6 with CM replaced by the letter-indicated (see text for definitions of E, S, D and D^*′*^).

| Measure<br>Approach | $\text{Eff}^\sigma$ | $\text{Eff}^\kappa$ | $\text{Eff}^{\kappa^*}$ | $D_{JS\%}^{\text{ens}}$ | $E^\kappa$ |
| --- | --- | --- | --- | --- | --- |
| *all values are percentages* |  |  |  |  |  |
| $\mathcal{T}_{\text{sim}}^{\text{loc}}$ | | | | | |
| $\mathcal{M}_{5,S_4;3,S_4}$ | 94 | 97 | 96 | 35 | 19 |
| $\mathcal{M}_{5,D_4;3,D_4}$ | 90 | 89 | 89 | 38 | 14 |
| $\mathcal{M}_{10,E_4;3,E_4}$ | 99 | 95 | 96 | 24 | 24 |
| $\mathcal{M}_{15,S_4;3,S_4}$ | 97 | 90 | 87 | 17 | 36 |
| $\mathcal{M}_{15,D_4;3,D_4}$ | 95 | 89 | 93 | 25 | 16 |
| $\mathcal{T}_{\text{owl}}^{\text{loc}}$ | | | | | |
| $\mathcal{M}_{5,S_4;3,S_4}$ | 99 | 88 | 81 | 6.7 | 48 |
| $\mathcal{M}_{15,E_4;3,E_4}$ | 81 | 61 | 44 | 7.7 | 39 |
| $\mathcal{M}_{10,S_8;5,S_4}$ | 86 | 68 | 69 | 28 | 10 |
| $\mathcal{M}_{10,D_8;5,D_4}$ | 94 | 73 | 76 | 34 | 8.6 |
| $\mathcal{M}_{10,D'_8;5,D'_4}$ | 94 | 77 | 81 | 34 | 7.4 |

Some insights can also be obtained by graphing the word distributions obtained in when clustering the words into raw CAMs, particularly if the words remaining in the raw clusters and those reassigned from the raw cluster are identified. Such graphs become cumbersome if the number of words is large (e.g. when *n* = 8 and *m* = 5, *n*^*m*^ = 32, 768). Thus we only provide such graphs of our results for the two baseline cases (simulation and owl data) for which *n* = 4, *m* = 3 and hence *n*^*m*^ = 64. (Fig 2)

## DISCUSSION

The results recorded in Table 3, indicate that one of the most challenging features of our approach is to identify if not the best method parameters and approaches to clustering then, at least, a set that performs best among those compared. Apart from the clustering techniques themselves, the parameter values to be selected are segmentation steps (*µ*), StaME cluster number (*n*), word coding size (*m*) and CAM cluster number (*k*) parameters. Of the three clustering methods (i.e., CM=E, S, and D) that we used to undertake the CAM extraction part of our method (i.e., Tasks 3-6, Box 1), the deep-learning shape method D outperformed the spectral vector method S with regard to providing the lowest E^*κ*^ values for both the *µ* = 5 and *µ* = 15 basic segment size cases. We did not include a comprehensive comparative evaluation of Ward’s vector method (E), once we noticed that both 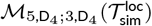 (Fig 2A, E^*κ*^ = 14%) and 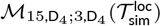 (Fig 2B, E^*κ*^ = 16%), outperformed our baseline method 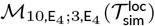 (Fig 2C, E^*κ*^ = 24%), which had a baseline segment size of *µ* = 10 intermediate between the two mentioned comparatives.

For all the simulation data 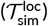 comparisons, we only considered the case *m* = 3 (i.e., CAMs consist of three StaMEs). In all cases where we compared coding efficiency in terms of segment size (i.e., *µ* = 5 versus *µ* = 15) for the same clustering methods used to extract both StaMEs and CAMs, higher error rates E^*κ*^ were obtained in the case of the smaller segment size (19% vs 36% in the case of S_4_ and 14% vs 16% in the case of D_4_ in Table 3). Additionally, the Jensen-Shannon divergence of the word distributions across the ensemble of clusters versus the original pool had small percentage divergences 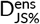 in these two same comparisons (35% versus 17% in the case of S_4_, and 38% vs 25% in the case of D_4_) for segment size. Thus, apart from a higher information density, smaller segment sizes appear to produce more reliable codes when the variety of StaME types is small (i.e., *n* around 4 rather than 8), provided segments are not too small to lose their shape diversity (e.g., only segments with three steps or more can exhibit squiggles).

The barn owl data presented a much greater challenge than the simulation data to find a set of parameters that provided CAM coding wtih low assignment error rates E^*κ*^. Our initial efforts in setting the variety of StaMEs to *n* = 4 and the number of StaMEs in a CAM to *m* = 3 produced error rates of E^*κ*^ = 48% and 39% respectively for the cases 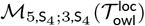 and 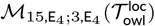 (Table 3). With considerable experimentation (results not included here), we were only able to reduce the error rates E^*κ*^ for the owl data to 10% or less by increasing the variety of StaMEs from *n* = 4 to 8 and the number of StaMEs used to represent a CAM from *m* = 3 to 5.

When we compare the StaME and rectified CAM statistics for the method 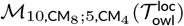 with CM=S, D and D^*′*^, the D method provided the greatest spread and the D^*′*^ method the smallest spread in velocity ranges across the four rectified CAMs (Table 1): a 13-fold spread for D (0.015-0.198), a 6.5-fold spread for S (0.020-0.131), and a 6-fold spread for D^*′*^ (0.018-0.107). The method CM=D^*′*^, however, provided a marginally more efficient coding (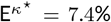, Table 3), than CM=D 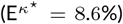 or S 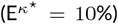. The distributions of StaMEs types in each of the rectified CAMs for these three cases are provided in Table 2. Not surprisingly, the method CM=D has its fastest CAM strongly dominated by its fastest StaME (when compared to the cases CM=D^′^ and S: 73.6% versus 41.9% and 42.8% respectively), given that we have already seen that CM=D has the greatest CAM velocity spread of the three.

We note from Table 1 that the primary discriminating factor among similarly valued StaMEs generated by a particular method is their net displacement (or “openness”). For example, for 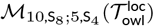, *σ*_6_, *σ*_7_ and *σ*_8_ all have 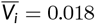, but their respective net-displacement values are 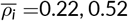 and 0.37. We should not expect this phenomenon to be as obvious for CAMs as it is for StaMEs because CAMs, being several times large than StaMEs, may be discriminated on shape features other than net displacement (e.g. number of zig-zags).

As an aside, we note that mixed clustering approaches—e.g., using S to cluster the StaMEs and D to cluster the CAM—appears to perform less well than using the same method for both clustering Tasks 2 and 4 in Box 1. For example the mixed approaches case 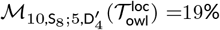 19%, (Table D2) has higher values when compared with the same approaches cases 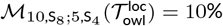 and 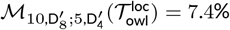. (Table 3).

In this paper we have presented a method for constructing a two-tier hierarchy of StaME and CAM building blocks underlying behavioral activity modes (BAMs). Our formulation can be thought of as bringing a microscope to bare on the fine structure of the statistical elements underlying the BAMs. Currently, BAMs are identified directly from a relocation data set *T* ^loc^ using BCPA (behavioral change point analysis ^13–17^), HMM (hidden Markov model ^18–22^) methods and clustering approaches ^9^ with no way to compare methods other than how well a particular BAM construction fits ground-truthed empirical data or the known structure of simulation data. On the other hand, the two-tiers of building blocks underlying variable length BAM segments of *T* ^loc^ in our approach allows us to employ information theoretic concepts to evaluate the efficiency of the StaME coding time series *T* ^*σ*^ (Box 1, Task 2) in terms of the extracted rectified CAM set *K*^⋆^ (Box 1, Task 6) in the absence of any ground truthing information. Of course, ground truthing can still be used in our approach to provide additional confirmation of how well our constructed BAMs fit empirical or simulated data.

How to use the information obtained from the various measures depicted in Table 3 is going to be an import problem to resolve in the future. High effective StaME and CAM entropies (Eff^*σ*^, 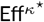, Table 3) are not necessarily indicators of good code because we should expect some movement modes to be rarer than others (e.g., hunting predators dashing at maximum speed versus cantering at sub-maximum speeds ^55^). The locations of where rare movement modes occur on the landscape (such as along human created seismic lines ^56^), as well as their relative frequencies and times of occurrence within the diel and seasonal cycles, are going to be important unraveling elements of the movement ecology of animals. It will likely take a number of studies using our methods on various types of organisms and environmental situations for a fuller appreciation of the utility and limitations of our information theoretic measures to emerge.

On the other hand, our measures now provide us with a method for testing hypotheses relating to the amount of information contained in particular animal movement tracks. Examples of such hypothesis may be that the information content of a movement track i) increases with maturity and experience of individuals, ii) decreases with stress and indicators of their poor health, iii) are correlated with particular individual personality traits ^43^, or iv) correlated with particular environmental and landscape variables. They may also be used to help us address questions of how animal movement behavior is responding to global change and to build predictive models of how such movement will adapt over time ^57^. At this time, most current real-world track data are too coarse to analyze using our segmentation methods, but in the new era of big data ^58^ this situation is rapidly changing. From the following calculations, it is obvious, that sub-minute data is required for our method to identify CAMs at a finer scale than 15 minute segments. For example, if relocation data is collected at 1 minute intervals and grouped into 5 min segments (i.e., *µ* = 5), with 3 segments constituting a word (i.e., *m* = 3), then this would at best provide a set of 15 min long CAM elements.

The barn owl data that we analyzed, though, is part of a high frequency (i.e., sub-minute) multi individual, multi-year barn owl data set collected in the Harod Valley using an ATLAS reverse GPS Atlas tracking system ^43^. For such data collected at a frequency of 0.25 Hz (4 sec interval), and with *µ* = 10 and *m* = 8 for the lowest three reassignment error values E^*κ*^ in Table 3, our CAM elements were 4 × 10 × 5 = 200 sec 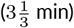 duration. The distributions of StaMEs in the rectified CAMs for these three cases indicated that none of the rectified CAMs came close to being behaviorally homogeneous (i.e., all their underlying StaMEs are of the same type). The closest we came to producing a homogeneous CAM was the case 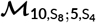 where we see in Table 2 that the percentage of fastest StaME (*σ*_1_) underling the fastest CAM 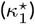 is 73.6% (implying that, on average, 2 of the 8 StaMEs instances of 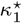 are not of type *σ*_1_). To obtain, at least one or more homogeneous Harod Valley barn owl rectified CAMs would likely require higher frequency data (e.g., 1 Hz), short StaMEs, fewer StaMEs per CAM, or a combination of all three.

## CONCLUSION

The approach presented here complements approaches that may begin by identifying different types of diel activity routines (DARs) ^40,59^ and then go on to carve these into subdiel behavioral activity modes (BAMs) using behavioral change point or hidden Markov models. Autoregressive time series models ^60^ can then be fitted to such BAMs to obtain a set of underlying canonical activity modes (CAMs). Our approach to identify CAMs from the”bottom up” rather than the before described “top down” approach allows us to impose an information theoretic structure on either simulated or fitted tracks at diel scales and above (e.g., seasonal life history movement phases or a complete life time track itself) ^31^. The primary limitation of our bottom up approach, however, is the existence of relatively high frequency movement data.

Future studies are needed to see how well our best sets of parameters and methods, as well as, other untested combinations of parameter values perform in providing new insights into the movement ecology of the Harod Valley barn owl or other populations for which high frequency data is available. For birds that typically fly at speeds considerably faster than the average movement speed of terrestrial animals, as well as move in three dimensions, it remains to be seen what time scale is needed to extract pure StaME-coded CAMs (perhaps data at the frequency of 1 Hertz are needed). Thus we might speculate that in the near future our methods are best tested on terrestrial rather than avian species.

A next step in applying our method to the Harod Valley barn owl data or any other high frequency relocation data should be to assess how the basic statistics of the extracted StaME and CAM elements depend on various factors, such as time of day or year, the sex or age of individuals, and variation across populations of individuals. Additionally, it will be particularly insightful to ascertain how StaME, CAM and BAM statistics are affected by underlying landscape variables. Of particular interest will be to incorporate accelerometer data, as it becomes increasingly available, into the specification and clustering of the base segments. This would be a powerful supplement to the analysis of high frequency location data itself, with our expectation that high frequency accelerometer and relocation data are likely to become increasingly available in the near future for all types of species ^58^, including terrestrial ^61^, arboreal ^62^, aerial ^63^, avian ^64^ and aquatic ^65^ species.

### Symbol Glossary

Parameters used to designate the numbers and proportions of movement track symbols, words, cluster types, and methods of analysis

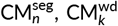 Refers to basic segment and word segment clustering methods with cluster number parameters *n* and *k* respectively (the five types used are CM = E, S, F, D and M)

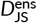 Jensen-Shannon ensemble divergence (Eq 5)

Eff^x^ x = *σ, κ* or *κ*^⋆^: efficiency of code x.

*E*^*κ*^ : raw to rectified CAM assigment error (Task 6, Box 1)

H^x^, 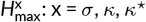 information entropies and max possible entropies

*k*: number of word clusters

*m*: number of symbols per word

*m*^*n*^ : number of word types *ω*_*l*_ ∈ *W* (not be confused with CAMs per se) 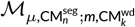 functor defined in Eq 6

*n*: number of basic segment clusters (StaME ideals *σ*_*i*_, *i* = 1, …, *n*)

*N*^seg^ = [*T*/*µ*]: number of basic segments in *T* ^seg^

*N*^wd^ = [*T*/(*mµ*)]: number of words in *T* ^wd^ in set *W*

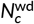: number of words in cluster *W*_*c*_, where 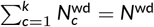

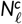: number of words of type *ω*_*l*_ ∈ *W*_*c*_, *l* = 1, …, *n*^*m*^

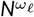: number of words of type *ω*_*l*_ ∈ *W*, *l* = 1, …, *n*^*m*^

**p**^*σ*^, 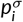: vector and proportions of symbols of type *σ*_*i*_ ∈ *S*

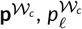: vector and proportions of words of type *l* in word cluster

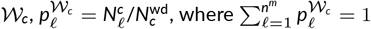

**p**^*κ*^, 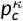 vector and proportions of all words in cluster *W*_*c*_ (prop. *κ*_*c*_) seg_*z*_, *z* = 1, …, *N* ^seg^ : Segments used to code *T* ^seg^

*T*: number points in the track excluding origen (point index *t* = 0, 1, …, *T*)

*T* ^loc^: The track as a relocation points time series

*T* ^*σ*^: The track as a StaME symbol time series

*T* ^wd^: The track as a word segment time series

*T* ^*κ*^: The track as a raw CAM time series

*T* ^CAM^: The track as a rectified CAM time series

*V*_*z*_,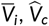: Average velocity of basic segment, StaME and CAM (vector representation Eq 9-11)
wd_*j*_: The *j*-th word in the time series *T* ^wd^

*W*: Set of words in track *T* ^wd^

*W*_*c*_ : Words in word segment cluster *c* (*c* = 1 …, *k*)

(*x*_*t*_, *y*_*t*_), *t* = 0, …, *T*: points in relocation time series *T* ^loc^

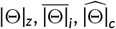: Average absolute turning angle of basic segment, StaME and CAM (vector representation Eq 9-11)

*κ*_*c*_, *K*: Raw CAMs *c* = 1, …, *k* and set thereof (Task 4, Box 1)

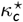, *K*^⋆^: rectified CAMs *c* = 1, …, *k* and set thereof (Task 6, Box 1)

*µ*: number of steps per basic segments

*ρ*, 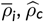: Average net displacement (distance between start of first end of last step) of basic segment, StaME and CAM (vector representation Eq 9-11)

*σ*_*i*_, *S*: StaMEs *i* = 1, …, *n*, and set of StaMEs

*ω*_*l*_, Ω: The *l*-th word type in the word type set Ω (Task 5, Box 1)

## Supporting information

(SOF)

Two_Kernel_Movement.csv

## Abbreviations

BCPA: behavioral change point analysis
HMM: hidden Markov methods
StaMEs: statistical movement elements
CAMs: canonical activity modes
BAMs: behavioral activity modes
ANIMOVER: ANImal MOVEment Ramp application’s simulator
SOF: supplementary online file

## Authors contribution

**VS** implemented all the clustering procedures, carried out all the numerical computations, contributed to drafting the manuscript. **OS** set up the project to collect the barn owl data and is a Co-PI of the project funding this study, made extensive comments on progressive versions of the ms. **RS** built the Numerus coding platform and RAMP simulator used to obtain the simulation data. **SC** collected and collated the barn owl data. **ST** is responsible for the technical and software aspects of the ATLAS system used to collect the barn owl data and is a Co-PI of the project funding this study. **WMG** designed this study, developed the theoretical underpinnings, with contributions from VS drafted the manuscript, and is a Co-PI of the project funding this study. **All** authors read and edited various versions of the manuscript including the final version

## Acknowledgments

Assaf Uzan, Michal Handel, Nadav Amir, Shani Cain and Yohay Wasserlauf helped with fieldwork and other components of data collection. We are also grateful to many people and institutes who permit us to use their towers and power supply for our base stations. These include Eli Chupa and Pablo Uner (Harod Valley Water Supply Co-op LTD), Shmuel Wirzberger (Kibbutz Beit Hashita), Eytan Ofer (Kibbutz Sde Eliyahu), Yitzhak Rivlin, Yado Shalev and Elon Shaked (Kibbutz Ein HaNetziv); Shaldag Refrigeration Storeroom LTD, Avihu Levanon (Partner PHI LTD), Michael Zelmanov (Pelephone Communications Ltd), Asher Nahmani and Rafi Levi (Mekorot), and Eli Cohen and Eli Mimran (Bezeq Inc.), and The Israeli Telecommunication Corp Ltd.

## DECLARATIONS

### Ethical Approval

Trapping and tagging procedures were authorized by permits 2019/42155 and 2020/42502 from Israel Nature and Parks Authority.

## Conflict of Interest

Wayne Getz and Richard Salter have a financial relationship with Numerus, Inc., a company that could benefit from the commercialization of the platform used to generate the simulation data analyzed in this research.

## Consent to participate

Not applicable.

## Consent to publish

Not applicable.

## Funding

This work was funded in by the Koret-UC Berkeley-Tel Aviv University Initiative in Computational Biology and Bioinformatics (WMG, OS, ST) and by a grant from the Data Science Center at Tel Aviv University (TAD). SC was also supported by the Hoopoe Foundation, Society for the Protection of Nature in Israel, Ministry of Agriculture and Rural Development, Ministry of Regional Cooperation, Larry Kornhauser, and Peter and Naomi Neustadter.

## Availability of data

The simulation data can be found in the supplementary online file (SOF) Two_Kernel_Movement.csv. The data used for the barn owl analysis is available at [LLV’s GitHub repository](https://github.com/LudovicaLV/DAR_project), as provided in ^40^

## Access to code

The code use to generate StaME and CAMs from relocation data and produce the various measures can be found at [VS’s GitHub repository](https://github.com/Observarun/statistical-movement-elements_canonical-activity-modes_CAM-coding-with-StaMEs).

## Supporting information

A supporting online file (SOF) containing Appendices A-D is provided with this document.

## Notes

https://github.com/Observarun/statistical-movement-elements_canonical-activity-modes_CAM-coding-with-StaMEs

https://github.com/LudovicaLV/DAR_project

### Summary of Updates

Minor editing and updating links to Github websites

## REFERENCES

1. McClintock BT, London JM, Cameron MF, Boveng PL. Bridging the gaps in animal movement: hidden behaviors and ecological relationships revealed by integrated data streams. Ecosphere. 2017;8(3):e01751.

2. Sih A, Spiegel O, Godfrey S, Leu S, Bull CM. Integrating social networks, animal personalities, movement ecology and parasites: a framework with examples from a lizard. Animal behaviour. 2018;136:195–205.

3. Kays R, Crofoot MC, Jetz W, Wikelski M. Terrestrial animal tracking as an eye on life and planet. Science. 2015;348(6240):aaa2478.

4. Allen AM, Singh NJ. Linking movement ecology with wildlife management and conservation. Frontiers in Ecology and Evolution. 2016;3:155.

5. Katzner TE, Arlettaz R. Evaluating contributions of recent tracking-based animal movement ecology to conservation management. Frontiers in Ecology and Evolution. 2020;7:519.

6. Edelhoff H, Signer J, Balkenhol N. Path segmentation for beginners: an overview of current methods for detecting changes in animal movement patterns. Movement ecology. 2016;4(1):21.

7. Smouse PE, Focardi S, Moorcroft PR, Kie JG, Forester JD, Morales JM. Stochastic modelling of animal movement. Philosophical Transactions of the Royal Society B: Biological Sciences. 2010;365(1550):2201–2211.

8. Spiegel O, Harel R, Centeno-Cuadros A, Hatzofe O, Getz WM, Nathan R. Moving beyond curve fitting: using complementary data to assess alternative explanations for long movements of three vulture species. The American Naturalist. 2015;185(2):E44–E54.

9. Teimouri M, Indahl U, Sickel H, Tveite H. Deriving Animal Movement Behaviors Using Movement Parameters Extracted from Location Data. ISPRS International Journal of Geo-Information. 2018 02;7:78.

10. Nathan R, Spiegel O, Fortmann-Roe S, Harel R, Wikelski M, Getz WM. Using tri-axial acceleration data to identify behavioral modes of free-ranging animals: general concepts and tools illustrated for griffon vultures. Journal of Experimental Biology. 2012;215(6):986–996.

11. Wilson AM, Hubel TY, Wilshin SD, Lowe JC, Lorenc M, Dewhirst OP, et al. Biomechanics of predator–prey arms race in lion, zebra, cheetah and impala. Nature. 2018;554(7691):183–188.

12. Barraquand F, Benhamou S. Animal movements in heterogeneous landscapes: identifying profitable places and homogeneous movement bouts. Ecology. 2008;89(12):3336–3348.

13. Chen J, Gupta AK. Parametric statistical change point analysis: with applications to genetics, medicine, and finance. Springer Science & Business Media; 2011.

14. Matteson DS, James NA. A nonparametric approach for multiple change point analysis of multivariate data. Journal of the American Statistical Association. 2014;109(505):334–345.

15. Gurarie E, Andrews RD, Laidre KL. A novel method for identifying behavioural changes in animal movement data. Ecology letters. 2009;12(5):395–408.

16. Gurarie E, Bracis C, Delgado M, Meckley TD, Kojola I, Wagner CM. What is the animal doing? Tools for exploring behavioural structure in animal movements. Journal of Animal Ecology. 2016;85(1):69–84.

17. Thompson PR, Harrington PD, Mallory CD, Lele SR, Bayne EM, Derocher AE, et al. Simultaneous estimation of the temporal and spatial extent of animal migration using step lengths and turning angles. Movement Ecology. 2024;12(1):1.

18. Franke A, Caelli T, Hudson RJ. Analysis of movements and behavior of caribou (Rangifer tarandus) using hidden Markov models. Ecological Modelling. 2004;173(2-3):259–270.

19. Langrock R, King R, Matthiopoulos J, Thomas L, Fortin D, Morales JM. Flexible and practical modeling of animal telemetry data: hidden Markov models and extensions. Ecology. 2012;93(11):2336–2342.

20. Michelot T, Langrock R, Patterson TA. moveHMM: An R package for the statistical modelling of animal movement data using hidden Markov models. Methods in Ecology and Evolution. 2016;7(11):1308–1315.

21. Zucchini W, MacDonald IL, Langrock R. Hidden Markov models for time series: an introduction using R. Chapman and Hall/CRC; 2016.

22. Pohle J, Langrock R, van Beest FM, Schmidt NM. Selecting the number of states in hidden Markov models: pragmatic solutions illustrated using animal movement. Journal of Agricultural, Biological and Environmental Statistics. 2017;22(3):270–293.

23. Fryxell JM, Hazell M, Börger L, Dalziel BD, Haydon DT, Morales JM, et al. Multiple movement modes by large herbivores at multiple spatiotemporal scales. Proceedings of the National academy of Sciences. 2008;105(49):19114–19119.

24. Benhamou S. Of scales and stationarity in animal movements. Ecology letters. 2014;17(3):261–272.

25. Sur M, Woodbridge B, Esque TC, Belthoff JR, Bloom PH, Fisher RN, et al. Linking behavioral states to landscape features for improved conservation management. Ecology and Evolution. 2021;11(12):7905–7916.

26. Owen-Smith N, Goodall V, Fatti P. Applying mixture models to derive activity states of large herbivores from movement rates obtained using GPS telemetry. Wildlife Research. 2012;39(5):452–462.

27. Owen-Smith N, Goodall V. Coping with savanna seasonality: comparative daily activity patterns of A frican ungulates as revealed by GPS telemetry. Journal of Zoology. 2014;293(3):181–191.

28. Gundermann KP, Diefenbach DR, Walter WD, Corondi A, Banfield J, Wallingford B, et al. Change-point models for identifying behavioral transitions in wild animals. Movement Ecology. 2023;11(1):65.

29. Getz WM, Salter R, Sethi V, Cain S, Spiegel O, Toledo S. The Statistical Building Blocks of Animal Movement Simulations. bioRxiv. 2023;p. 2023–12.

30. Getz WM, Saltz D. A framework for generating and analyzing movement paths on ecological landscapes. Proceedings of the National Academy of Sciences. 2008;105(49):19066–19071.

31. Getz WM. A hierarchical path-segmentation movement ecology framework. Ecological Processes. 2022;11(1):1–15.

32. Morales JM, Haydon DT, Frair J, Holsinger KE, Fryxell JM. Extracting more out of relocation data: building movement models as mixtures of random walks. Ecology. 2004;85(9):2436–2445.

33. Austin D, Bowen W, McMillan J. Intraspecific variation in movement patterns: modeling individual behaviour in a large marine predator. Oikos. 2004;105(1):15–30.

34. Benhamou S. Detecting an orientation component in animal paths when the preferred direction is individual-dependent. Ecology. 2006;87(2):518–528.

35. Van Moorter B, Visscher DR, Jerde CL, Frair JL, Merrill EH. Identifying Movement States From Location Data Using Cluster Analysis. The Journal of Wildlife Management. 2010;74(3):588– 594.

36. Jaeger A, Banks D. Cluster analysis: A modern statistical review. Wiley Interdisciplinary Reviews: Computational Statistics. 2023;15(3):e1597.

37. Tarca AL, Carey VJ, Chen Xw, Romero R, Drăghici S. Machine learning and its applications to biology. PLoS computational biology. 2007;3(6):e116.

38. Thessen A. Adoption of machine learning techniques in ecology and earth science. One Ecosystem. 2016;1:e8621.

39. Xie J, Girshick R, Farhadi A. Unsupervised deep embedding for clustering analysis. In: International conference on machine learning. PMLR; 2016. p. 478–487.

40. Luisa Vissat L, Cain S, Toledo S, Spiegel O, Getz WM. Categorizing the geometry of animal diel movement patterns with examples from high-resolution barn owl tracking. Movement Ecology. 2023;11(1):1–20.

41. Cagnacci F, Focardi S. Animal Movement. In: Renso C, Spaccapietra S, Zimányi E, editors. Mobility Data Modeling, Management, and Understanding. Cambridge University Press; 2013. p. 259 – 276.

42. Getz WM. An Information Theory Treatment of Animal Movement Tracks. In: Giuggioli L, Maini P, editors. The Mathematics of Movement: An Interdisciplinary Approach to Mutual Challenges in Animal Ecology and Cell Biology. New York: Springer; 2024. p. 2403.16290.

43. Cain S, Solomon T, Leshem Y, Toledo S, Arnon E, Roulin A, et al. Movement predictability of individual barn owls facilitates estimation of home range size and survival. Movement Ecology. 2023;11(1):10.

44. Manning C, Schutze H. Foundations of statistical natural language processing. MIT press; 1999.

45. Ren Y, Pu J, Yang Z, Xu J, Li G, Pu X, et al. Deep clustering: A comprehensive survey. 221004142. 2022;.

46. Lafabregue B, Weber J, Gançarski P, Forestier G. End-to-end deep representation learning for time series clustering: a comparative study. Data Mining and Knowledge Discovery. 2022;36(1):29–81.

47. Shannon CE. A mathematical theory of communication. The Bell system technical journal. 1948;27:379–423.

48. Shannon CE. A mathematical theory of communication. The Bell system technical journal. 1948;27:623–656.

49. Toledo S, Kishon O, Orchan Y, Shohat A, Nathan R. Lessons and experiences from the design, implementation, and deployment of a wildlife tracking system. In: Software Science, Technology and Engineering (SWSTE), 2016 IEEE International Conference on. IEEE; 2016. p. 51–60.

50. Beardsworth C, Gobbens E, van Maarseveen F, Denissen B, Dekinga A, Nathan R, et al. Validating ATLAS: a regional-scale, high-throughput tracking system. Methods Molecular Evolution. 2022;.

51. Cuturi M, Blondel M. Soft-dtw: a differentiable loss function for time-series. In: International conference on machine learning; 2017..

52. von Luxburg U. A tutorial on spectral clustering. Statistics and Computing. 2007;17:395–416.

53. Parmar A, Katariya R, Patel V. A review on random forest: An ensemble classifier. In: International conference on intelligent data communication technologies and internet of things (ICICI) 2018. Springer; 2019. p. 758–763.

54. Ratanamahatana CA, Keogh E. Everything you know about Dynamic Time Warping is Wrong. In: Third Workshop on Mining Temporal and Sequential Data, in conjunction with the Tenth ACM SIGKDD International Conference on Knowledge Discovery and Data Mining (KDD-2004); 2004..

55. Wilson AM, Lowe J, Roskilly K, Hudson PE, Golabek K, McNutt J. Locomotion dynamics of hunting in wild cheetahs. Nature. 2013;498(7453):185–189.

56. McKenzie HW, Merrill EH, Spiteri RJ, Lewis MA. How linear features alter predator movement and the functional response. Interface focus. 2012;2(2):205–216.

57. Getz WM, Luisa Vissat L, Salter R. Simulation and analysis of animal movement paths using Numerus Model Builder. In: 2020 Spring Simulation Conference (SpringSim). IEEE; 2020. p. 1–12.

58. Nathan R, Monk CT, Arlinghaus R, Adam T, Alós J, Assaf M, et al. Big-data approaches lead to an increased understanding of the ecology of animal movement. Science. 2022;375(6582):eabg1780.

59. Owen-Smith N. Daily movement responses by African savanna ungulates as an indicator of seasonal and annual food stress. Wildlife Research. 2013;40(3):232–240.

60. Biswas A, Guha A. Time series analysis of categorical data using auto-mutual information. Journal of Statistical Planning and Inference. 2009;139(9):3076–3087.

61. Papageorgiou D, Farine DR. Group size and composition influence collective movement in a highly social terrestrial bird. Elife. 2020;9:e59902.

62. Borah B, Beckman NG. Studying seed dispersal through the lens of movement ecology. Oikos. 2022;2022(2).

63. Niga Y, Fujioka E, Heim O, Nomi A, Fukui D, Hiryu S. A glimpse into the foraging and movement behaviour of Nyctalus aviator; a complementary study by acoustic recording and GPS tracking. Royal Society Open Science. 2023;10(6):230035.

64. Conners MG, Michelot T, Heywood EI, Orben RA, Phillips RA, Vyssotski AL, et al. Hidden Markov models identify major movement modes in accelerometer and magnetometer data from four albatross species. Movement ecology. 2021;9:1–16.

65. Francisco FA, Nührenberg P, Jordan A. High-resolution, noninvasive animal tracking and reconstruction of local environment in aquatic ecosystems. Movement ecology. 2020;8:1–12.

