## Supplementary material for "An Information Theory Framework for Movement Path Segmentation and Analysis": (SOF)

### Supplementary online file of Appendices to:
### Sethi et al., An Information Theory Framework for Movement Path Segmentation and Analysis

#### A DATA PREPARATION

As discussed in [1], we set the time parameters in the ANIMOVER_1 RAMP to produce $100$ time steps for each of our "nominal days". The variable `Delta`, which is equivalent to *t*, resets at the turn of each day. This mandates a transformation on this variable since we need it to be continuous (sans jumps from $99$ to $0$) for segmentation. We find the average speed and turning angle across the steps of a single segment. Following the computation of a reflection invariant absolute turning angle $|\Delta\theta(t)|$ [1,2], the very first point is discarded for lack of a value at $|\Delta\theta(0)|$. Speeds above $70$ meters per second are discarded for empirical data.

After completing the above, we now have an ordered sequence of the relevant variables. For further calculations, a normalization is performed on speed by scaling values using the maximum observed value over all points in the complete trajectory, while turning angle is scaled by $\pi$. Consequently, we have $v(t) \in [0,1]$, $\Delta\theta(t) \in (-1,1]$, and $|\Delta\theta(t)| \in [0,1]$. The normalized series is then parsed into a set of base segments $\mathcal{T}^{\rm seg}$ (or $\mathcal{T}^{\rm wd}$) each with $\mu$ (or $m\mu$) steps to construct a representation required by vector or shape clustering methods, as described in the next sub-section. For owl data, segmentation is performed through time variable (`POSIXct` type in `R`) while enforcing the constraint of $\mu$ (or $m\mu$) points in each segment. This lets us handle missing points in that the segments with fewer than $\mu$ (or $m\mu$) points drop out. A clustering procedure is then performed on the symbolic sequence $\mathcal{T}^{\sigma}$ (or word sequence $\mathcal{T}^{\kappa}$) to construct the set $\mathcal{S}$ of StaMEs as shown in Task 2 of Box 1 (or the set $\mathcal{K}$ of CAMs, Tasks 3, 4).

#### B CLUSTERING METHODS

As mentioned in the main text, the methods we used to cluster the data can be divided into two classes: vector methods that use euclidean and machine learning approaches [3–7], and shape methods that use deep learning approaches [8–10]. Some details of these methods are provided below.

##### Vector Methods

Vector based methods are applied to the statistical characteristic vectors of each segment, as represented by Eq 10 and 11 in the main text. The following vector clustering methods (CM) were used in our analyses.

**CM=E** Hierarchical clustering is applied to the Euclidean (E) representation (E; Eq 1 and 2 in the main text) using Ward's criterion [7] and a dissimilarity matrix constructed using a Euclidean measure. The dissimilarity matrix is computed using `R 4.3.2:parallelDist::parDist()`, while clustering has been performed using `R:fastcluster::hclust()`. This method was used in the context of Eq 6 (main text) to obtain 8-StaMEs sets for the data analyzed here [1].

**CM=S** Clustering is performed on the data transformed into similarity space as follows. Graph spectral (S) clustering [11] first constructs a similarity graph from the data using the affinity matrix: a matrix of point-to-point similarities $A_{ij} = exp(-\frac{||x_i - x_j||^2}{2\sigma^2})$ (Gaussian kernel with Euclidean metric), where points form the nodes while edges quantify the similarity between the points. Clusters are then found using k-means method in the reduced dimensional representation extracted from the spectral analysis of the normalized graph Laplacian $L = \mathbb{1} - D^{-1/2} A D^{-1/2}$, where diagonal node degree matrix $D_{ii} = \sum_{j=1}^{n} A_{ij}$. Summarily, it's a k-way graph cut problem—the connections between the points with low similarity are severed, while the net weight of the edges between the clusters are minimized. The computational complexity of the algorithm is $\mathcal{O}(N_{seg}^3)$. We implemented this method in `Python 3.10.10` using a Gaussian radial base function (RBF) kernel

**CM=F** Random Forest (F) classifiers [12] can be used in an unsupervised way to generate a proximity matrix. The algorithm creates a synthetic dataset with each variable randomly sampled from the univariate distribution of the corresponding variable of the original dataset. The original and new datasets are given class labels $1$ and $2$, and merged to be used for training the random forest. The proximity matrix, constructed in terms of how often two points end up in the same leaf nodes, is then used for hierarchical clustering with Ward's criterion using `R:randomForest::randomForest` with `proximity` variable set to `TRUE`.

##### Shape methods

Shape-based clustering methods are applied directly to segments of the time-series data $\mathcal{T}^{v}$ (Eq 7 in main text) and $\mathcal{T}^{\Delta\theta}$ (Eq 8 in main text) or, as the case may be, the absolute values). The following methods were used in our analyses.

**CM=D** and **D′** Dynamic Time Warping [13] (DTW) is a temporal shift-invariant similarity measure that performs one-to-one and one-to-many maps (or temporal alignments) between two time-series to minimize the Euclidean distance between aligned series. The difference between D and D′ is that the former uses absolute turning angle data, while the latter uses actual turning angle data.

The algorithm constructs a local cost matrix (LCM) employing an $L_p$-norm ($p \in \{1,2\}$) between each pair of points in the two time-sequences (lengths $m$, $n$), with the $ij^{\rm th}$ element given by $D(i,j) = d(i,j) + min\{d(i-1,j); d(i,j-1), d(i-1,j-1)\}$. Finding an optimal warp path then amounts to minimizing the cost over all admissible paths from LCM$(1,1)$ to LCM$(n,m)$ while ensuring that the sequence of index pairs increases monotonically in both $i$ and $j$. We use the R `dtw_basic` function to help manage the $\mathcal{O}(N_{seg}^2)$ computational complexity with its C++ core and memory optimizations.

To ensure the warping path remains close to the diagonal, the slanted band window constraint is used. The `stepPattern` employed to traverse through the LCM is R `symmetric2`, which is commonly used and makes our DTW computation symmetric. The clustering solution selected for different choices of window size within the range $\{1, \ldots, 4\}$ and value of $p = 1$ or 2 in $L_p$-norm selected was the one that either provided the largest silhouette coefficient value (a value of 1 implies perfect separation of clusters, though the largest value we obtained for owl data was usually < $0.1$). In case the solution with the second largest coefficient value had a much better average velocity spread across the cluster centroids, we selected that solution based on that fact that larger velocity spreads provide a better bases for discrimination among movement modes (e.g. running versus striding versus ambling, etc.).

**CM=M** DTW cost is a non-differentiable function, which limits its utility when used as a loss function. Soft-min-DTW[14] is a differentiable alternative to DTW making use of a soft-min (M) operator.

#### C NUMBERING SYMBOLS AND WORDS

The material presented here, augments the discussion on this topic provided in the main text.

As an example of our numbering scheme, consider a segmentation where the number of symbols is 3 ($n = 3$) with $V_1 > V_2 > V_3$ and words are two symbols long. Then, using the notation $\mathbf{V}^{\mathrm{wd}} = V_{i_1} + \cdots + V_{i_m}$ for words made up of the $m$ symbols represented by the indices $\{i_1, \cdots, i_m$, it follows that if we number the words using a sorting algorithm that prioritizes the first index over the second when word sizes are equal then our word ordering will be

$$\begin{aligned} &\omega_1 \sim \sigma_1\sigma_1\sigma_1,\ \omega_2 \sim \sigma_1\sigma_1\sigma_2,\ \omega_3 \sim \sigma_1\sigma_2\sigma_1,\ \omega_4 \sim \sigma_2\sigma_1\sigma_1, \\ &\omega_5 \sim \sigma_1\sigma_2\sigma_2,\ \omega_6 \sim \sigma_2\sigma_1\sigma_2,\ \omega_7 \sim \sigma_2\sigma_2\sigma_1,\ \text{and}\ \omega_8 \sim \sigma_2\sigma_2\sigma_2 \\ &\Rightarrow \{\omega_1, \cdots, \omega_8\} \equiv \{111, 112, 121, 211, 122, 212, 221, 222\} \end{aligned} \tag{C1}$$

In the addition the velocity relationships will be

$$\mathbf{V}^{\omega_1} > \mathbf{V}^{\omega_2} = \mathbf{V}^{\omega_3} = \mathbf{V}^{\omega_4} > \mathbf{V}^{\omega_5} = \mathbf{V}^{\omega_6} = \mathbf{V}^{\omega_7} > \mathbf{V}^{\omega_8} \tag{C2}$$

The case $n = 4$, $m = 3$ is provided in Eq 12 in the main text. Other cases for $n \leq 3$ follow using the same numbering convention of grouping words that add up to the same symbol index value together, starting with the largest of the indices on the right hand side. For the case $n = 5$, 6 and 8, we use the expected quinary, heximal, and octal numbering systems for 3-digit numbers (case $m = 3$), except we start at 1 rather than 0 and fill in all three digits (1-5, 1-6, and 1-8 respectively). In this case, however, words of equal size are not grouped together, though $\omega_1 \equiv 11..1$ is still the largest word and $\omega_{n^m} \equiv n..n$ is still the smallest word. Thus, for the case $n = 5$ and $m = 3$ the numbering process is

$$\begin{aligned} \{\omega_1, \cdots, \omega_{125}\} \equiv \big\{&111, 112, 113, 114, 115, 121, 122, 123, 124, 125, 211, \\ &212, 213, 214, 215, 311, \cdots, 553, 554, 555\big\} \end{aligned} \tag{C3}$$

and similarly for $n = 6$ and $8$, and $m = 3$ and 4, as the case may be.

### D ADDITIONAL TABLES

Additional tables referred to in the text as residing in the Appendix are presented in this section

**TABLE** D1

Comparison of the mean step length (velocity) $\overline{V}_i$ and mean absolute turning angles $\overline{|\Delta\Theta|}_i$ of two (**A.**) and four StaME (**B.**) cluster centers (Eq 10, main text) for relocation data $\mathcal{T}_{\mathrm{sim}}^{\mathrm{seg}}$ base segments of size $\mu = 5$, 10, 15 and 30, clustered using E (Euclidean Wards), S (spectral), F (random forest), D (dynamic time warping) and M (soft-min dynamic time warping) methods discussed in the text. The efficiency **Eff** of each method is computed using Eq 4 (main text)

| | **Clustering on** seg$_z$ **vector representations (Eq 9, main text)** | | | | | | | | | **Clustering on** seg$_z$ $\mu$**-step time series** | | | | | |
|---|---|---|---|---|---|---|---|---|---|---|---|---|---|---|---|
| | **E** | | | **S** | | | **F** | | | **D** | | | **M** | | |
| ***Seg. size*** | | | **Eff** $^\sigma$ | | | **Eff** $^\sigma$ | | | **Eff** $^\sigma$ | | | **Eff** $^\sigma$ | | | **Eff** $^\sigma$ |
| $i$ | $\overline{V}_i$ | $\overline{\|\Delta\Theta\|}_i$ | %Tot. | $\overline{V}_i$ | $\overline{\|\Delta\Theta\|}_i$ | %Tot. | $\overline{V}_i$ | $\overline{\|\Delta\Theta\|}_i$ | %Tot. | $\overline{V}_i$ | $\overline{\|\Delta\Theta\|}_i$ | %Tot. | $\overline{V}_i$ | $\overline{\|\Delta\Theta\|}_i$ | %Tot. |
| **A.** | $n = 2$ **StaMEs** | | | | | | | | | | | | | | |
| ***5 step*** | | | **0.99** | | | **0.93** | | | **0.59** | | | **0.98** | | | **0.97** |
| 1 | 0.72 | 0.22 | 55% | 0.87 | 0.16 | 35% | 0.95 | 0.08 | 14% | 0.82 | 0.20 | 42% | 0.81 | 0.19 | 39% |
| 2 | 0.11 | 0.34 | 45% | 0.22 | 0.34 | 65% | 0.36 | 0.31 | 86% | 0.18 | 0.34 | 58% | 0.21 | 0.33 | 61% |
| ***10 step*** | | | **0.78** | | | **0.92** | | | **0.49** | | | **0.94** | | | **0.99** |
| 1 | 0.85 | 0.13 | 23% | 0.79 | 0.16 | 34% | 0.95 | 0.08 | 11% | 0.82 | 0.18 | 35% | 0.66 | 0.23 | 55% |
| 2 | 0.33 | 0.32 | 77% | 0.27 | 0.33 | 66% | 0.39 | 0.30 | 89% | 0.25 | 0.33 | 65% | 0.18 | 0.34 | 45% |
| ***15 step*** | | | **1.00** | | | **0.97** | | | **0.38** | | | **0.85** | | | **0.97** |
| 1 | 0.58 | 0.22 | 47% | 0.69 | 0.19 | 40% | 0.95 | 0.08 | 7% | 0.84 | 0.18 | 28% | 0.67 | 0.23 | 40% |
| 2 | 0.33 | 0.33 | 53% | 0.28 | 0.33 | 60% | 0.41 | 0.29 | 93% | 0.30 | 0.31 | 72% | 0.30 | 0.31 | 60% |
| ***30 step*** | | | **0.98** | | | **1.00** | | | **0.50** | | | **0.91** | | | **0.97** |
| 1 | 0.58 | 0.24 | 59% | 0.56 | 0.23 | 51% | 0.82 | 0.13 | 11% | 0.70 | 0.21 | 33% | 0.57 | 0.25 | 39% |
| 2 | 0.26 | 0.33 | 41% | 0.33 | 0.32 | 49% | 0.40 | 0.29 | 89% | 0.32 | 0.30 | 67% | 0.37 | 0.29 | 61% |
| **B.** | $n = 4$ **StaMEs** | | | | | | | | | | | | | | |
| ***5 step*** | | | **0.93** | | | **0.94** | | | **0.69** | | | **0.90** | | | **0.91** |
| 1 | 0.95 | 0.09 | 18% | 0.95 | 0.09 | 18% | 0.95 | 0.08 | 15% | 0.92 | 0.16 | 27% | 0.92 | 0.16 | 27% |
| 2 | 0.73 | 0.31 | 18% | 0.69 | 0.31 | 19% | 0.43 | 0.31 | 69% | 0.55 | 0.25 | 11% | 0.54 | 0.27 | 12% |
| 3 | 0.50 | 0.26 | 19% | 0.47 | 0.27 | 20% | 0.10 | 0.33 | 7% | 0.50 | 0.30 | 16% | 0.50 | 0.30 | 16% |
| 4 | 0.11 | 0.34 | 45% | 0.11 | 0.34 | 43% | 0.07 | 0.32 | 10% | 0.12 | 0.34 | 46% | 0.11 | 0.34 | 45% |
| ***10 step*** | | | **0.99** | | | **0.97** | | | **0.55** | | | **0.91** | | | **0.93** |
| 1 | 0.85 | 0.13 | 23% | 0.94 | 0.07 | 14% | 0.95 | 0.08 | 11% | 0.91 | 0.16 | 23% | 0.95 | 0.13 | 16% |
| 2 | 0.59 | 0.31 | 22% | 0.58 | 0.31 | 26% | 0.43 | 0.29 | 78% | 0.68 | 0.20 | 8% | 0.55 | 0.28 | 23% |
| 3 | 0.38 | 0.29 | 23% | 0.47 | 0.26 | 26% | 0.09 | 0.36 | 5% | 0.44 | 0.30 | 27% | 0.54 | 0.24 | 16% |
| 4 | 0.11 | 0.35 | 32% | 0.12 | 0.35 | 34% | 0.07 | 0.32 | 6% | 0.16 | 0.34 | 42% | 0.18 | 0.34 | 45% |
| ***15 step*** | | | **0.95** | | | **0.97** | | | **0.70** | | | **0.89** | | | **0.94** |
| 1 | 0.83 | 0.14 | 21% | 0.93 | 0.10 | 13% | 0.95 | 0.08 | 7% | 0.87 | 0.17 | 24% | 0.94 | 0.15 | 12% |
| 2 | 0.71 | 0.30 | 38% | 0.51 | 0.31 | 31% | 0.83 | 0.12 | 6% | 0.53 | 0.25 | 13% | 0.56 | 0.26 | 28% |
| 3 | 0.37 | 0.29 | 26% | 0.47 | 0.26 | 29% | 0.46 | 0.29 | 66% | 0.50 | 0.28 | 15% | 0.48 | 0.26 | 20% |
| 4 | 0.19 | 0.34 | 39% | 0.12 | 0.34 | 27% | 0.11 | 0.35 | 21% | 0.20 | 0.33 | 49% | 0.21 | 0.33 | 40% |
| ***30 step*** | | | **0.95** | | | **0.93** | | | **0.84** | | | **0.92** | | | **0.96** |
| 1 | 0.77 | 0.22 | 19% | 0.90 | 0.15 | 9% | 0.82 | 0.13 | 11% | 0.87 | 0.17 | 12% | 0.71 | 0.22 | 15% |
| 2 | 0.48 | 0.25 | 40% | 0.51 | 0.29 | 34% | 0.54 | 0.22 | 12% | 0.54 | 0.26 | 18% | 0.49 | 0.28 | 24% |
| 3 | 0.39 | 0.31 | 16% | 0.45 | 0.26 | 35% | 0.47 | 0.29 | 54% | 0.52 | 0.26 | 25% | 0.48 | 0.26 | 23% |
| 4 | 0.18 | 0.34 | 25% | 0.17 | 0.34 | 22% | 0.17 | 0.34 | 23% | 0.25 | 0.32 | 44% | 0.30 | 0.31 | 37% |

**TABLE** D2

For $x = \sigma$, $\kappa$ and $\kappa^\star$ the percent efficiencies $\mathrm{Eff}^x$ (Eq 4, main text) are listed for mixed clustering approaches (different method used to cluster StaMEs versus CAMs) along with the percent normalized Jensen-Shannon ensemble divergences $D^{\mathrm{ens}}_{\mathrm{JS\%}}$ (Eq 5, main text) and the percent assignment errors $\mathrm{E}^\kappa$ (Task 6, Box 1). The notation used is that of Eq 6 (main text) with CM replaced by the S, D and D′, as defined in Appendix B above.

| Measure / Approach | $\mathrm{Eff}^\sigma$ | $\mathrm{Eff}^\kappa$ | $\mathrm{Eff}^{\kappa^\star}$ | $D^{\mathrm{ens}}_{\mathrm{JS\%}}$ | $\mathrm{E}^\kappa$ |
|---|---|---|---|---|---|
| | *all values are percentages* | | | | |
| $\mathcal{M}_{5,\mathrm{S}_4;3,\mathrm{D}_4}(\mathcal{T}^{\mathrm{loc}}_{\mathrm{sim}})$ | 94 | 95 | 95 | 35 | 15 |
| $\mathcal{M}_{5,\mathrm{D}_4;3,\mathrm{S}_4}(\mathcal{T}^{\mathrm{loc}}_{\mathrm{sim}})$ | 90 | 97 | 98 | 33 | 28 |
| $\mathcal{M}_{15,\mathrm{S}_4;3,\mathrm{D}_4}(\mathcal{T}^{\mathrm{loc}}_{\mathrm{sim}})$ | 97 | 95 | 75 | 18 | 32 |
| $\mathcal{M}_{15,\mathrm{D}_4;3,\mathrm{S}_4}(\mathcal{T}^{\mathrm{loc}}_{\mathrm{sim}})$ | 95 | 90 | 90 | 18 | 32 |
| $\mathcal{M}_{10,\mathrm{S}_8;5,\mathrm{D}'_4}(\mathcal{T}^{\mathrm{loc}}_{\mathrm{owl}})$ | 86 | 77 | 78 | 27 | 19 |
| $\mathcal{M}_{10,\mathrm{D}'_8;5,\mathrm{S}_4}(\mathcal{T}^{\mathrm{loc}}_{\mathrm{owl}})$ | 94 | 68 | 70 | 32 | 7.9 |

#### REFERENCES


1. Getz WM, Salter R, Sethi V, Cain S, Spiegel O, Toledo S. The Statistical Building Blocks of Animal Movement Simulations. bioRxiv. 2023;p. 2023–12.
2. Getz WM. A hierarchical path-segmentation movement ecology framework. Ecological Processes. 2022;11(1):1–15.
3. Tarca AL, Carey VJ, Chen Xw, Romero R, Drăghici S. Machine learning and its applications to biology. PLoS computational biology. 2007;3(6):e116.
4. Van Moorter B, Visscher DR, Jerde CL, Frair JL, Merrill EH. Identifying Movement States From Location Data Using Cluster Analysis. The Journal of Wildlife Management. 2010;74(3):588–594.
5. Thessen A. Adoption of machine learning techniques in ecology and earth science. One Ecosystem. 2016;1:e8621.
6. Teimouri M, Indahl U, Sickel H, Tveite H. Deriving Animal Movement Behaviors Using Movement Parameters Extracted from Location Data. ISPRS International Journal of Geo-Information. 2018 02;7:78.
7. Jaeger A, Banks D. Cluster analysis: A modern statistical review. Wiley Interdisciplinary Reviews: Computational Statistics. 2023;15(3):e1597.
8. Xie J, Girshick R, Farhadi A. Unsupervised deep embedding for clustering analysis. In: International conference on machine learning. PMLR; 2016. p. 478–487.
9. Ren Y, Pu J, Yang Z, Xu J, Li G, Pu X, et al. Deep clustering: A comprehensive survey. arXiv:221004142. 2022;.
10. Lafabregue B, Weber J, Gançarski P, Forestier G. End-to-end deep representation learning for time series clustering: a comparative study. Data Mining and Knowledge Discovery. 2022;36(1):29–81.
11. von Luxburg U. A tutorial on spectral clustering. Statistics and Computing. 2007;17:395–416.
12. Parmar A, Katariya R, Patel V. A review on random forest: An ensemble classifier. In: International conference on intelligent data communication technologies and internet of things (ICICI) 2018. Springer; 2019. p. 758–763.
13. Ratanamahatana CA, Keogh E. Everything you know about Dynamic Time Warping is Wrong. In: Third Workshop on Mining Temporal and Sequential Data, in conjunction with the Tenth ACM SIGKDD International Conference on Knowledge Discovery and Data Mining (KDD-2004); 2004. .
14. Cuturi M, Blondel M. Soft-dtw: a differentiable loss function for time-series. In: International conference on machine learning; 2017. .
